# Iron Supplementation Eliminates Antagonistic Interactions Between Root Associated Bacteria

**DOI:** 10.1101/2020.06.05.132704

**Authors:** Thomas Eng, Robin A. Herbert, Uriel Martinez, Brenda Wang, Joseph Chen, Ben Brown, Adam Deutschbauer, Mina Bissell, Jenny C. Mortimer, Aindrila Mukhopadhyay

**Affiliations:** Joint BioEnergy Institute, Emeryville, CA, USA; Biological Systems and Engineering Division, Lawrence Berkeley National Laboratory, Berkeley, CA, USA; Environmental Genomics and Systems Biology Division, Lawrence Berkeley National Laboratory, Berkeley, CA, USA; Computational Biosciences Group, Computational Research Division, Computing Sciences Area, Lawrence Berkeley National Laboratory, CA, USA; Department of Statistics, University of California, Berkeley, Berkeley, CA, USA; Department of Plant & Microbial Biology, University of California, Berkeley, Berkeley, USA; Machine Learning and AI Group, Arva Intelligence Inc, Park City, UT, USA; College of Science and Engineering, San Francisco State University, San Francisco, California, USA

## Abstract

The rhizosphere microbiome (rhizobiome) plays a critical role in plant health and development. However the processes by which the constituent microbes interact to form and maintain a community are not well understood. To investigate these molecular processes, we examined pairwise interactions between 11 different microbial isolates under selected nutrient-rich and nutrient-limited conditions. We observed that when grown with media supplemented with 56 mM glucose, 2 microbial isolates were able to inhibit the growth of 6 out of 11 other microbes tested. The interaction between microbes persisted even after the antagonistic microbe was removed, upon exposure to spent media. To probe the genetic basis for these antagonistic interactions, we used a barcoded transposon library in a proxy bacterium, *Pseudomonas putida*, to identify genes which showed enhanced sensitivity to the antagonistic factor(s) secreted by *Acinetobacter* sp. 02. Iron metabolism-related gene clusters in *P. putida* were implicated by this systems-level analysis. The supplementation of iron prevented the antagonistic interaction in the original microbial pair supporting the hypothesis that iron limitation drives antagonistic microbial interactions between rhizobionts. We conclude that rhizobiome community composition is influenced by competition for limiting nutrients with implications for growth and development of the plant.

## Introduction

Microbial communities are being increasingly appreciated for their impact on larger biological systems, such as in relation to human health (e.g. the gut microbiome) (Chu et al., 2016) or crop productivity (e.g. the rhizobiome) (Mendes et al., 2018). It is now understood that antagonism between microbes can influence host fitness in various systems (Zhao et al., 2018; Mendes et al., 2018). However, while some direct chemical interactions between hosts and their microbiome are known (Stringlis et al., 2018), it is unclear how beneficial microbial communities are assembled and maintained throughout host development. A better understanding of microbial communities and their relation to the plant root is of specific interest to understand plant development, biogeochemical carbon cycling, and applications in agriculture.

The environmental milieu of the plant root is distinct from that of bulk soil. Photosynthetic bioproducts are exuded through the roots, modifying the rhizosphere through the accumulation of sugars and secondary plant metabolites. The composition of root exudate is complex (Chaparro et al., 2013; Kawasaki et al., 2016); however simple sugars (such as D-glucose) have often been detected as major components of the root exudate in terrestrial plants such as the model dicot *Arabidopsis thaliana* (Okubo et al., 2016). Changes in exudate profile correlate with compositional and functional changes in the rhizobiome (Chaparro et al., 2013).

While many isolates have been identified across ecologically distinct rhizobiomes, isolates from several major phyla tend to dominate any given rhizobiome (Vorholt, 2012; Bai et al., 2015). The formation of the rhizobiome is likely a deterministic process, in which microbes both synergize and antagonize each other in competition for limited resources from root exudate (Hassani et al., 2018). A population equilibrium, where several microbial phyla dominate, is eventually established. For example, specific microbes could be growth limited by the plant through the secretion of iron sequestering siderophores or antibiotics (Joshi et al., 2006; van der Meij et al., 2017). Alternatively, microbial inhibitory mechanisms such as secretion systems capable of puncturing neighboring cell membranes, or the secretion of antimicrobial compounds, have also been described (Alteri and Mobley, 2016; Nester, 2014). However, the specific environmental cues that prime a microbial response to limit the growth of other microbes remain poorly described.

In this study, we sought to understand if microbes from a model rhizobiome compete with each other under defined glucose supplementation conditions. We chose 11 rhizobacteria (**Table 1, Supplemental Figure 1**) representative of phyla previously detected as enriched within the roots of *A. thaliana* relative to bulk soil (Lundberg et al., 2012; Levy et al., 2017). These microbes are of general interest because they can improve plant health in the presence of specific environmental stressors (Herbert et al., 2019). We hypothesized that microbe-microbe interactions could be readily detected by tracking the frequency by which microbes inhibited the growth of their neighboring species. Out of 37 pairwise microbial competition assays in this format, we detected 3 microbes which either blocked colony formation of many microbes or specific microbial isolates. To examine the molecular mechanism underlying an inhibitory interaction, we used a pooled transposon mutant library (Wetmore et al., 2015) built in the soil microbe *Pseudomonas putida* KT2440 to identify genes required for growth in *Acinetobacter sp*. 02 spent medium, and validated our findings back in the synthetic rhizobiome microbe pair. A summary of our experimental approach is described in Figure 1.

**Figure 1.**
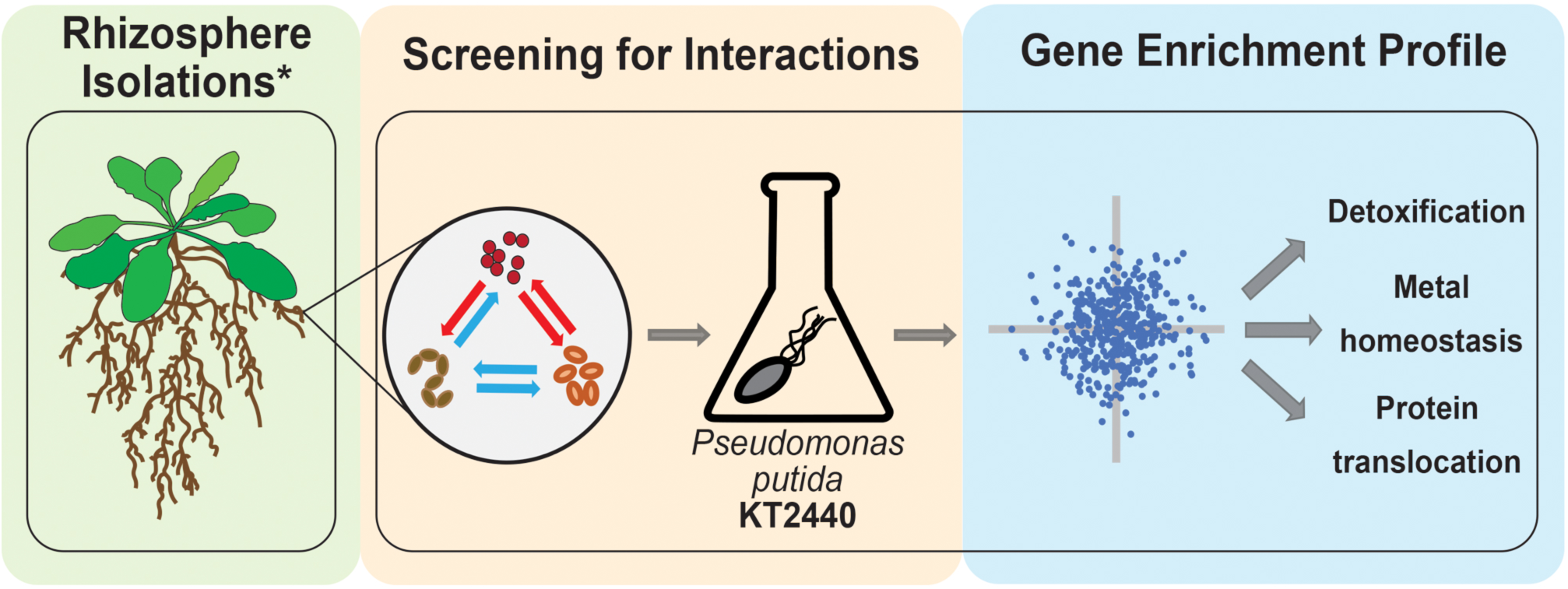
Workflow for bacterial chemical genomic screen. Rhizobacteria mediate complex interactions in the rhizosphere (green-shaded box). After isolation, rhizobacteria are screened for potential interactions and an RB-TnSeq library of a model microorganism (orange-shaded box). Finally, RB-TnSeq data are analyzed and used to characterize and validate microbe-microbe interactions.

## Results

### A concurrent inoculation screen to determine microbe-microbe interactions

We developed a rapid and reproducible assay to monitor microbe-microbe interactions based on cell viability as an alternative to technically involved methods, such as metagenomic RNAseq or microbial capture in microdroplets (Mutha et al., 2019; Jackman et al., 2019). We adapted an established agar plate based assay used to monitor cell-cell interactions in *Saccharomyces cerevisiae* (Abelson et al., 1990; Liu et al., 2019). When two microbial isolates are grown together **(Figure 2A)**, it should be possible to observe one of four types of interactions based on growth. First, there could be no detectable interaction between the two microbes. Second, both strains could still form colonies, but exhibit morphological changes or exhibit synergistic improved growth. Third, one of the microbes could fail to grow in the presence of the other microbe. Finally, both microbes could be growth-inhibited when co-cultivated. Bioinformatic analysis of these microbial genomes (Weber et al., 2015) to identify putative secreted gene clusters indicated that many of these species had the capacity to produce a range of secreted molecules, but it was unclear what conditions would be needed to induce production, or if any given microbe would be sensitive to a given secreted molecule **(Supplemental Table 1)**. A representative agar plate is shown in **Figure 2B**, where *Acinetobacter* sp. 02 is the primary species and tested for interactions with six secondary species. Finally, we modified this method to allow if the continued presence of cells was required for an antagonistic interaction with a staggered plating regime (see Materials and Methods).

**Figure 2.**
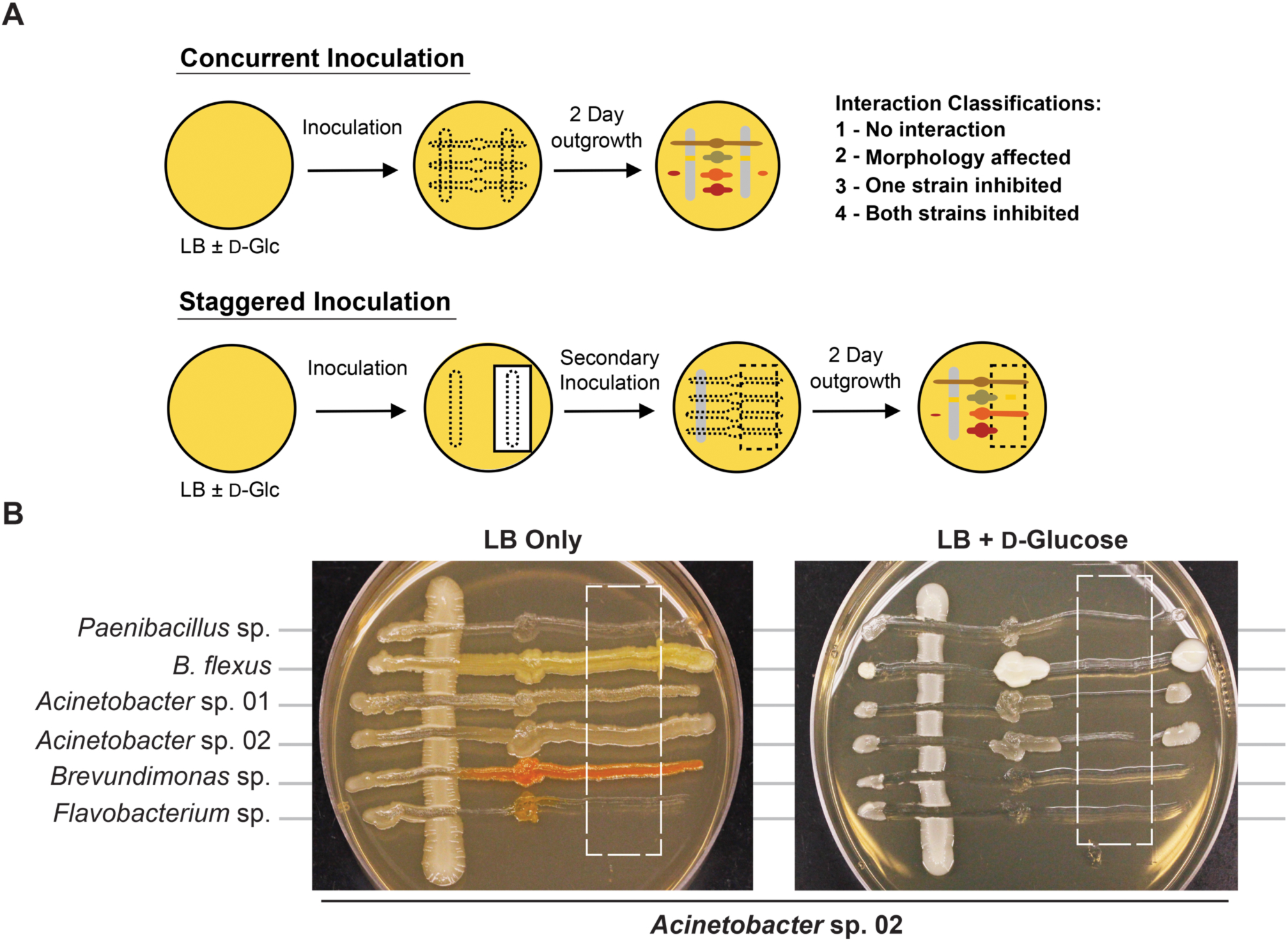
Methodology for microbial interaction screen. (A) Multiple rhizobacteria are used to inoculate LB plates, either concurrently or staggered such that the vertical, “primary” species was grown for 2 days before the introduction of a secondary species. For staggered inoculation assays, half of the primary inoculum was grown on top of a nitrocellulose membrane that was removed prior to secondary inoculation. (B) Representative images of staggered inoculation assays with *Acinetobacter* sp. 02 grown in co-culture with six rhizobacterial species. Dashed lines indicate the location of a 0.44 µm nitrocellulose membrane prior to removal. Brightness and contrast have been uniformly edited to increase visibility.

When the primary and secondary microbes were inoculated at the same time, we detected two pairwise interaction types. The majority of microbes tested for interactions showed no detectable change in growth when co-cultured **(Figure 3A)**. The following six isolates were screened for interactions (*A. rhizogenes, Acinetobacter* sp. 02, *Arthrobacter* sp., *Flavobacterium* sp., *Paenibacillus* sp., and *Ralstonia* sp.) against seven others (*A. rhizogenes, Acinetobacter* sp. 01, *Acinetobacter* sp. 02, *Bacillus flexus, Brevundimonas* sp., *Flavobacterium* sp., and *Paenibacillus* sp.; n ≥ 3) on solid LB (“limited glucose”). However, both *Acinetobacter* sp. 02 and *Flavobacterium* sp. specifically inhibited the growth of *Brevundimonas* sp. under the concurrent growth regimen **(Figure 3A)**.

**Figure 3.**
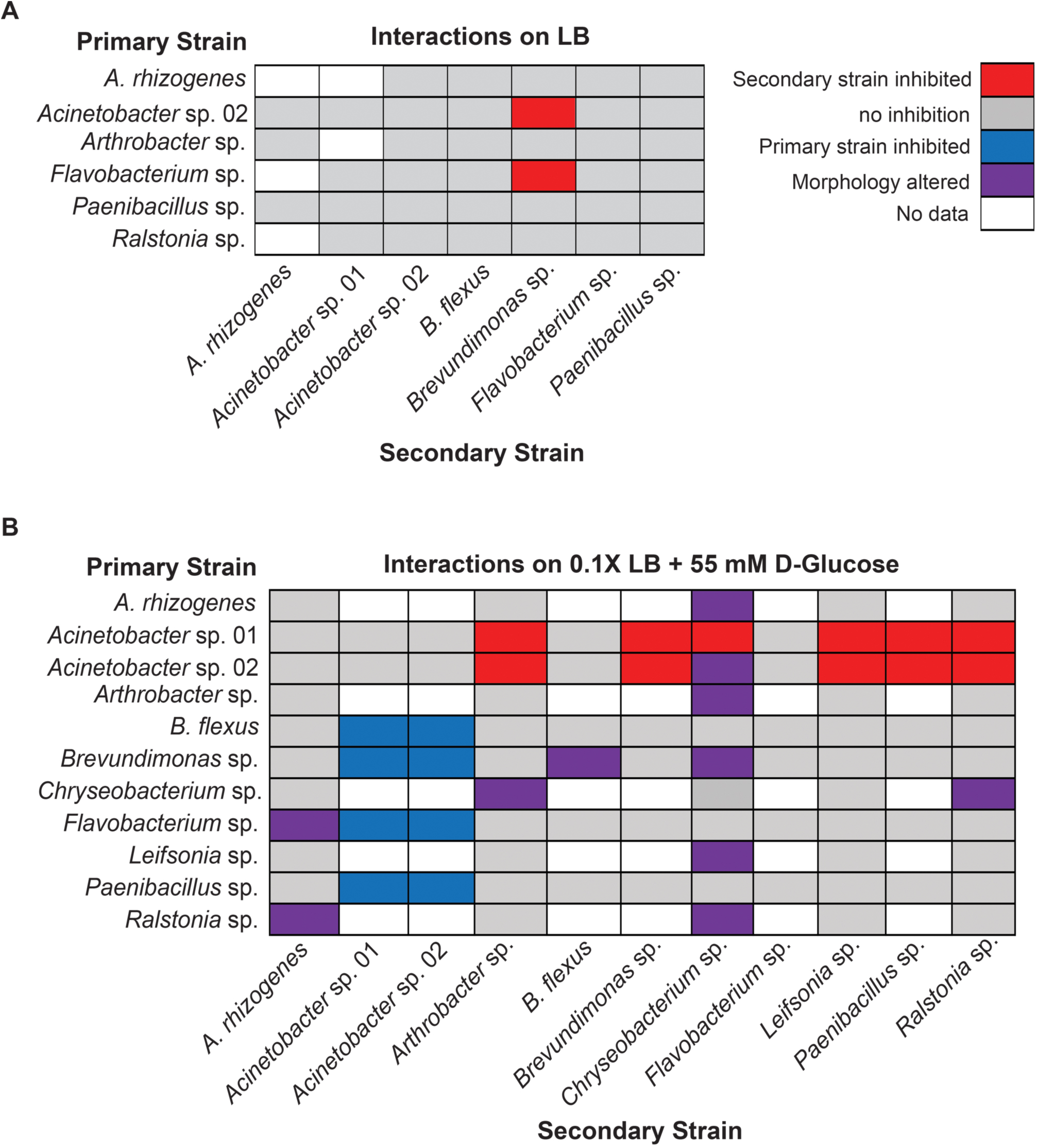
Results of concurrent inoculation interaction screen. Interactions observed on (A) LB or (B) 0.1X LB supplemented with 55 mM D-Glucose. Rows represent primary, vertically-streaked species and columns represent secondary, horizontally-streaked species.

Increasing the concentration of available glucose to 55 mM in the media allowed us to detect several additional cases of antagonistic interactions. We examined a larger set of pairwise microbial interactions (*Acinetobacter* sp. 01, *Acinetobacter* sp. 02, *Bacillus flexus, Brevundimonas* sp., *Flavobacterium* sp., and *Paenibacillus* sp., n ≥ 3 replicates) against the same set and added the following microbes: *A. rhizogenes, Arthrobacter* sp., *Chryseobacterium* sp., *Leifsonia* sp., and *Ralstonia* sp.; n ≥ 2).

Under the glucose supplemented media conditions additional inhibitory interactions were observed. Specifically, *Acinetobacter* sp. 01 inhibited the growth of seven different microbes: *Arthrobacter* sp., *Brevundimonas* sp., *Chryseobacterium* sp., *Flavobacterium* sp., *Leifsonia* sp., *Paenibacillus* sp., and *Ralstonia* s*p.* **(Figure 3B)**. *Acinetobacter* sp. 02 inhibited a similar set of microbes to *Acinetobacter* sp. 01 **(Figure 3B)**, with the exception of *Chryseobacterium* sp. In contrast, *Flavobacterium* sp., which was able to inhibit the growth of *Brevundimonas* sp. When grown in glucose supplemented conditions **(Figure 3A)**, had no effect when cultivated on LB media **(Figure 3B)**. However there were a number of interactions, e.g. between *B. flexus* and either of the *Acinetobacter* spp., wherein the order of inoculation could have a deterministic effect.

We also observed a range of morphological changes in the concurrent inoculation assay under glucose supplemented conditions. Examples include *Chryseobacterium* sp. changing colony morphology and color in the co-culture assay when grown with 6 of the 11 isolates tested **(Figure 3B, Supplemental Figure 2)**. *Ralstonia* sp. appeared to rapidly overtake cells of both *A. rhizogenes* or *Chryseobacterium* sp., but not other species such as *Arthrobacter* sp. or *Leifsonia* sp., consistent with our definition of a non-inhibitory microbial interaction **(Supplemental Figure 3)**. *Bacillus flexus* also appeared to grow more rapidly along *Brevundimonas* sp. (n=4 out of 6). Finally, *Flavobacterium* sp. formed an irregular border when co-cultured with *A. rhizogenes* **(Supplemental Figure 3)**. Together, these results suggest a variety of interactions may occur between different rhizobacterial pairs and the identification of three of the four types of potential microbial interactions.

### A staggered microbe inoculation screen identifies founder effects

Having identified three microbes with clear growth inhibitory phenotypes in our concurrent inoculation assay (*Flavobacterium* sp. and two isolates of *Acinetobacter* spp.), we next sought to understand the mechanism underlying these interactions. Of the three, both *Flavobacterium* sp. and *Acinetobacter* sp. 02 were able to inhibit growth of other microbes at a distance. We hypothesized that these strains could be secreting an environment-modifying molecule (e.g. an antimicrobial compound) or depleting a nutrient. This is in contrast to microbial type VI secretion systems, which requires direct cell contact to lyse competitors (Alteri and Mobley, 2016). In addition, while we did not detect any cell-cell interactions using *Paenibacillus* sp. in the concurrent inoculation assay **(Figure 3**), evidence from the literature suggested that members of the genus can produce growth inhibitory antimicrobials (Shaheen et al., 2011; Meng et al., 2018). Therefore, we chose these three microbes to test whether they could alter the growth media such that it was not inhibitory to other microbes.

We first used the staggered inoculation regime to test for microbial interactions. *Acinetobacter* sp. 02, *Flavobacterium* sp., and *Paenibacillus* sp. were grown for two days either on a sterile nitrocellulose membrane placed on top of the solid agar media. After two days, the membrane and cells were removed, resulting in spent solid media. Alternatively, the microbes were plated directly onto solid media (refer to **Figure 2** for diagram and representative plate). The second microbe was then streaked onto the plate (**Figure 2**). Being able to directly assess growth conditions on the same agar plate strengthens our ability to detect microbial interactions in spent agar media. *Flavobacterium* sp. and *Paenibacillus* sp. grew poorly on the nitrocellulose membrane under nutrient-poor conditions (data not shown), and so standard LB media was used for these assays instead. Following removal of *Acinetobacter* sp. 02, *Flavobacterium* sp. was still inhibited **(Figure 4A)**, the only case in which this antagonism of *Flavobacterium* sp. was observed. As a primary strain, *Flavobacterium* sp. pretreatment inhibited the growth of both *Brevundimonas* sp., as well as freshly plated *Flavobacterium* sp. itself, however this antagonism not was observed on the spent solid media **(Figure 4A)**. Similarly, *Paenibacillus* sp. was inhibitory against *Bacillus flexus* in the presence of *Paenibacillus* sp. cells only **(Figure 4A)**. No other interactions between other microbe pairs in the staggered assay were detected.

**Figure 4.**
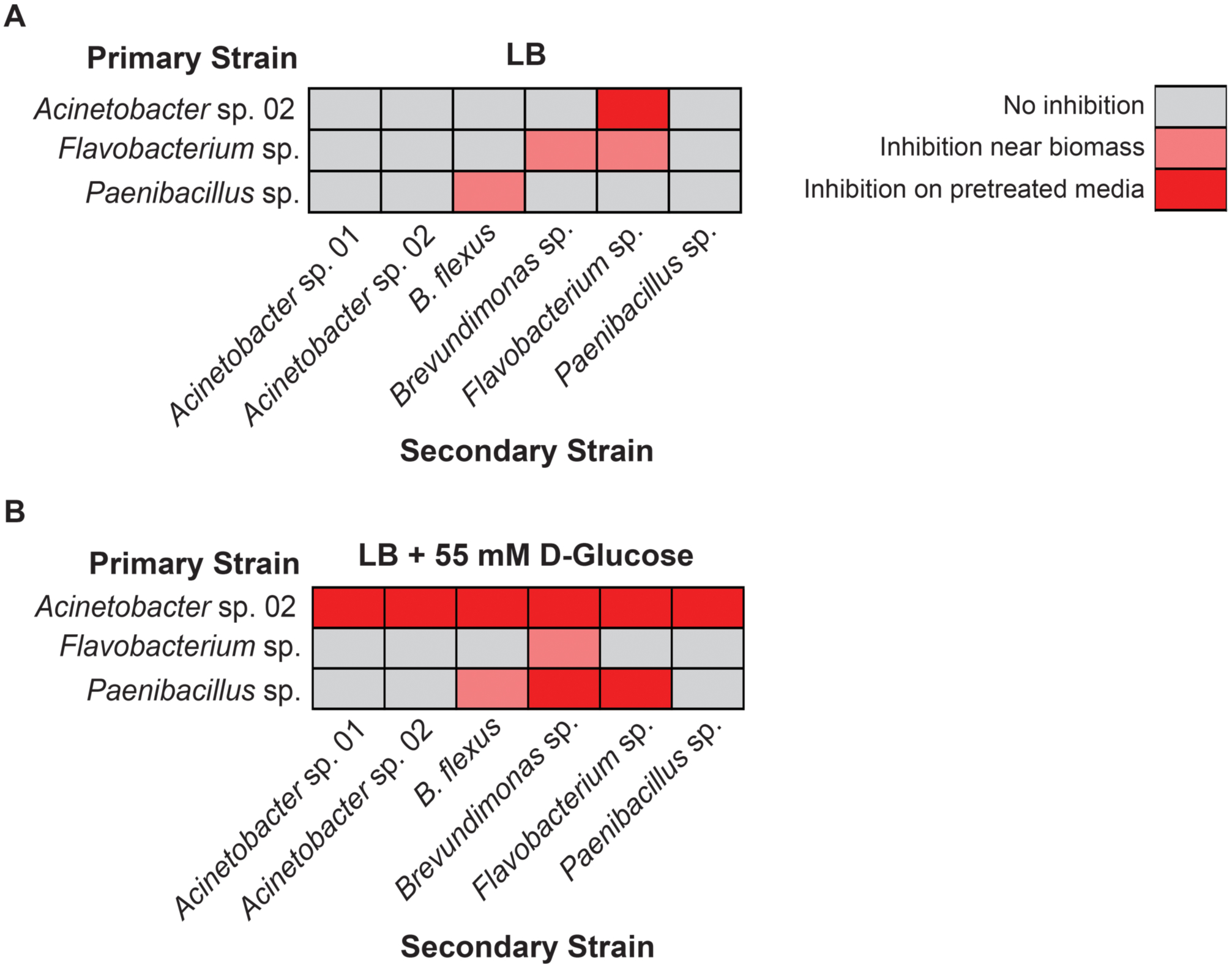
Results of staggered inoculation interaction screen. Interactions observed on (A) LB or (B) LB supplemented with 55 mM D-Glucose. Rows represent primary, vertically-streaked species and columns represent secondary, horizontally-streaked species. n ≥ 3. Where interactions were variable by colony, the most frequently observed interaction is displayed.

Next, we repeated the staggered inoculation assay with glucose supplemented media to determine if the spent solid media was now more or less toxic. We observed that in this experiment, *Acinetobacter* sp. 02 was inhibitory to all six tested microbes both with and without the cells present (see representative plate in **Figure 2B, Figure 4B)**. *Flavobacterium* sp. inhibited the growth of *Brevundimonas* sp. and no other **(Figure 4B)**. Finally, spent solid media from *Paenibacillus* sp. was inhibitory to both *Brevundimonas* sp. and *Flavobacterium* sp. **(Figure 4B)**. This data suggests that specific microbial interactions do not require direct cell-cell contact, and can persist even after a microbe is removed from the environment. These interactions are differentially mediated by D-glucose concentration/ availability. These observations support our hypothesis that growth inhibition arises through depletion of an essential nutrient or the secretion of an antimicrobial compound.

### *Using RB-TnSeq in P. putida to identify secreted factors from Acinetobacter* sp. 02

Having established that *Acinetobacter* sp. 02 modified its environment such that it was inhibitory to a range of different microbes, we sought to identify the secreted factor(s) that conferred this growth inhibitory phenotype. Since genetic tools are not yet available for these recently isolated microbes, we used *Pseudomonas putida* KT2440 as a proxy. *P. putida* KT2440 is an established soil microbe found in similar environments as many microbes in our representative rhizobiome (Molina, 2000). Wild-type *P. putida* KT2440 is not sensitive to the *Acinetobacter* sp. 02 supernatant (**Supplemental Figure 4**). Accordingly, we hypothesized that individual genes in *P. putida* KT2440 which are responsive to the potential antagonistic agent/ condition present in the media would provide insight into the phenotype being observed in the rhizobacterial pairs. Using a barcoded transposon library for parallel fitness assays is an ideal method for the rapid identification of such mutants (Wetmore et al., 2015), which has been generated for *P. putida* KT2440. This barcoded library contains ∼100,000 unique transposon mutants with coverage of most nonessential genes (Thompson et al., 2019; Rand et al., 2017). By growing these mutants in a pooled format, *P. putida* mutants which are sensitive to the toxic agents in the supernatant will be outcompeted by more fit strains, and the absolute abundance of each mutant can be determined using Illumina sequencing specific to each barcoded transposon mutant. Analysis of transposon abundances are used to implicate gene functions that are correlated with resistance/ susceptibility to the secreted molecule (**Figure 5A)**.

**Figure 5.**
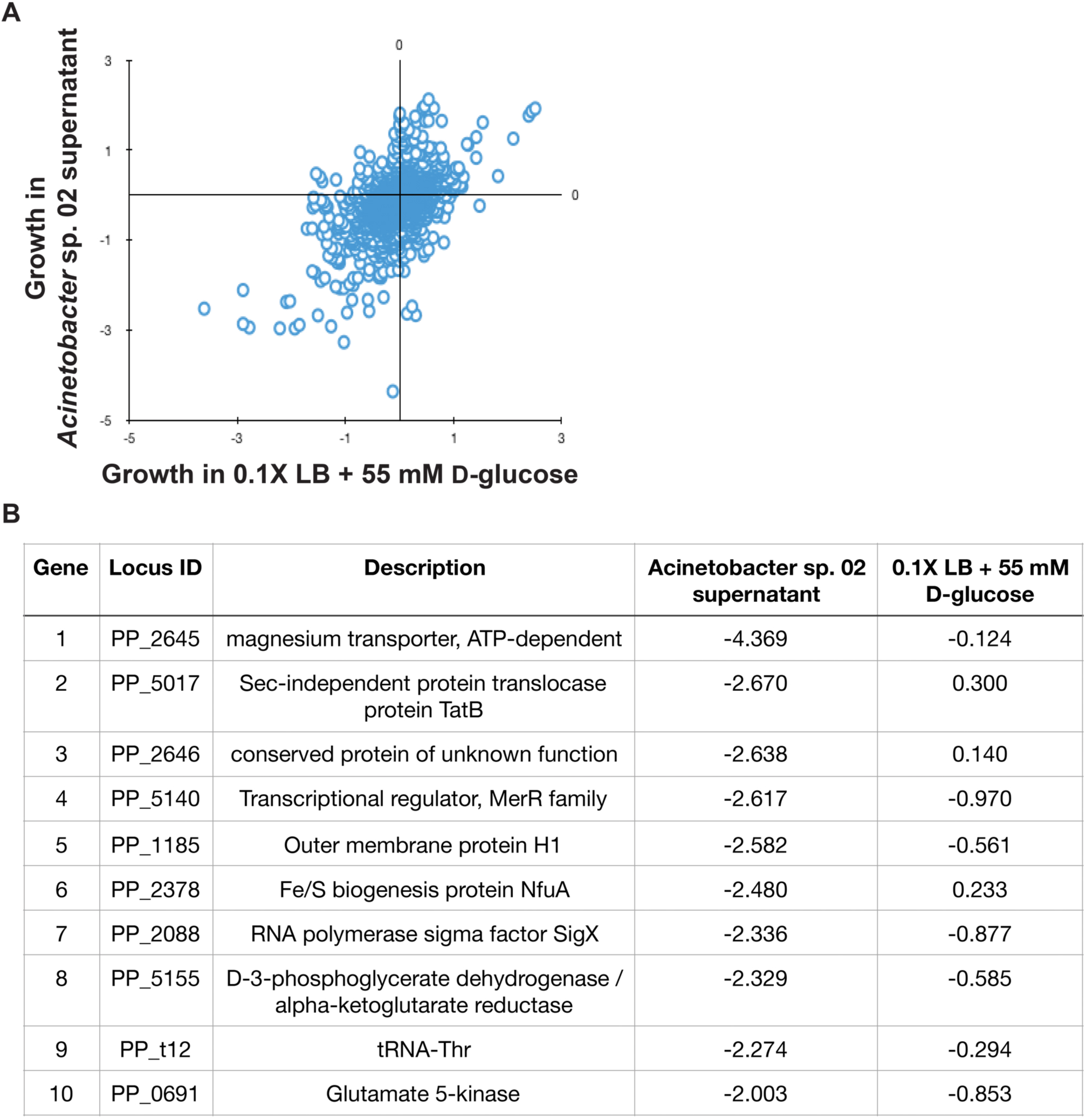
*Pseudomonas putida* KT2440 RB-TnSeq results. (A) Scatterplot of gene fitness values from *P. putida* KT2440 grown in *Acinetobacter* sp. 02 supernatant vs. control media. (B) The top ten depleted genes in library populations grown in supernatant and relevant log_2_-fold decreases relative to the population at Time 0.

In the RB-TnSeq data, 10 genes had a statistically significant fitness defect, relative to the control conditions. Interestingly, many of these genes with phenotypes are involved in metal ion transport or metabolism. Both genes in a two gene operon, *PP_2645* and *PP_2646*, were sensitive to the *Acinetobacter* sp. 02 supernatant. *PP_2645* shows sequence similarity to ATP-dependent magnesium transporters **(Figure 5B)** while *PP_2646* remains uncharacterized **(Figure 5B)**. Mutants in the metal responsive transcriptional regulator, *PP_5140* (*merR*, (Miller et al., 2009)) and a metal-responsive outer membrane protein *PP_1185* were also sensitive to this supernatant **(Figure 5B)**. Moreover, we also recovered mutants in *PP_2378*, a candidate Fe/S related protein *(NfuA)* **(Figure 5B)**. A comparison with an additional biological replicate of the control condition also implicated two other metal ion related genes, *PP_3244* and *PP_5139*, which may be related to magnesium and cadmium transport. Both full, gene-for-gene comparisons can be found in Supplementary Data Set 1. Together, these gene targets suggested that the *Acinetobacter* sp. 02 supernatant was growth-limiting due to the absence or inactivation of an essential metal cofactor such as magnesium or iron.

### Validation of iron requirement in synthetic rhizobiome pair

Next, we tested whether *Acinetobacter* sp. 02 could be reducing iron availability in the media, for example by sequestering metal ions with a siderophore like pyoverdine (Drehe et al., 2018). Elemental iron is a limiting and essential nutrient in all cells (Cairo et al., 2006). Our bioinformatics analysis (**Supplemental Table 1**) indicated that several of the representative microbes had the capacity to produce siderophores, and *Acinetobacter* sp. 02 encodes a putative siderophore that was weakly similar (22% identity) to vicibactin (Wright et al., 2013) (**Supplemental Table 1**). We tested this hypothesis by repeating our concurrent inoculation assay with *Acinetobacter* sp.02 and several secondary strains, but examined conditions where cells were grown in nutrient-rich, dilute glucose medium with or without supplemental iron. LB media contains approximately 9.6 µM iron from yeast peptone and tryptone. Consistent with our predictions from our functional genomics analysis, iron supplementation of the media restored growth of *Brevundimonas* sp. and *Leifsonia* sp. **(Figure 6A and 6B)**. The effect of iron supplementation was dose-dependent, as 10 and 100 µM FeCl3 were sufficient to improve *Brevundimonas* sp. and *Leifsonia* sp. growth in the presence of *Acinetobacter* sp. 02, but 1 µM FeCl3 was not **(Supplemental Figure 5)**.

**Figure 6.**
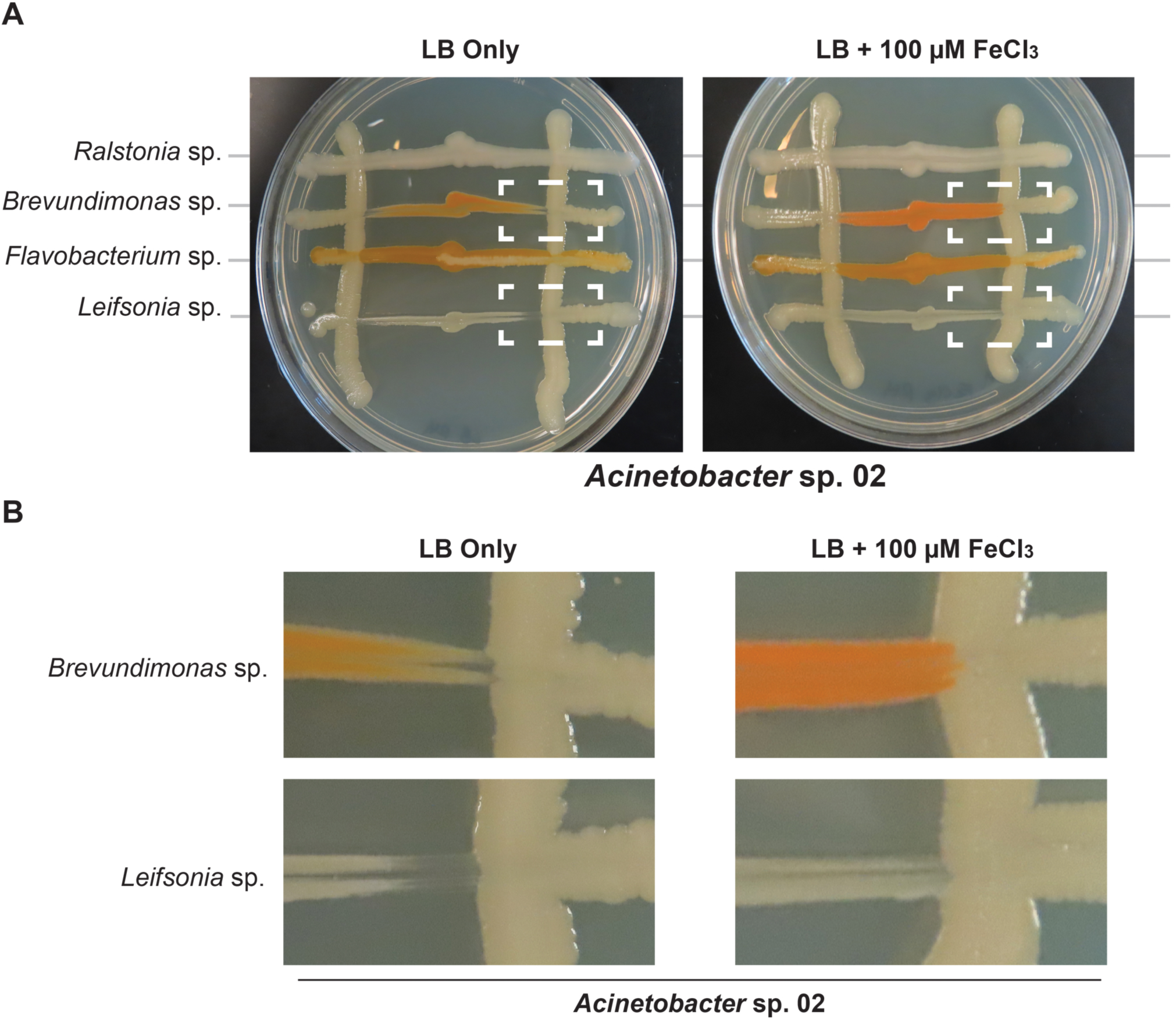
Iron supplementation in *Acinetobacter* sp. 02 concurrent interaction assay. *Acinetobacter* colonies were streaked vertically on LB with or without 100 µM supplemental FeCl_3_ followed by the perpendicular inoculation of the secondary species indicated. Dashed lines indicate interactions that change depending on iron supplementation. n ≥ 3.

## Discussion

Microbes interact with the root and each other to play an important role in the architecture of the root-associated microbial communities and their function. However, the relationship between the root and its associated microbes is complex and remains to be disentangled. In this report we dissect the pairwise interactions which may occur between several representative bacterial members of the *Arabidopsis* root microbiome. We tested predictions generated from analyzing the behavior of a proxy soil microbe in response to a complex spent growth media using a systems biology approach. This analysis generated predictive insights on the potential growth inhibitory molecule in the *Acinetobacter* sp. 02 supernatant, which in turn helped our understanding of microbial interactions between two genetically unmodified rhizobacterial species. From this analysis, we provide evidence that competition for iron in the culture media can physiological changes in *Acinetobacter* sp. 02 such that it acts to limit competition from other microbes in its vicinity. Our work sheds light on the context by which individual microbes may respond to the specialized nutrient cues from the nutritional environment near the root, as the broad growth inhibitory activity of *Acinetobacter* sp. 02 is dependent on specific nutrient conditions. Moreover, the founder effects we observed in this study could confound higher degree microbial interaction studies if the order of species arrival in an environment is not carefully controlled.

Competition for essential metals beyond iron represents a general mechanism likely to affect rhizobiome assembly. Both plant and animal hosts have been observed to engage in nutritional immunity, wherein hosts sequester essential metals in order to limit the growth of potential opportunistic pathogens. Therefore, the ability of these bacteria to compete for metals, both in relation to their host and other microbes, could be general mechanisms that influence host colonization. For instance, iron and zinc homeostasis are essential for colonization of mammalian host tissues by the pathogen *Acinetobacter baumannii (Lonergan et al., 2019; Hood et al., 2012)*. Similarly, root-associated *Acinetobacter* sp. have been observed to sequester metals such as copper (Rojas-Tapias et al., 2012, 2014), thereby affecting plant growth. In our system, we deduced that iron was a key driver of microbial antagonism; iron has been shown to be a key factor in shaping both plant exudates and the microbiome (Butaitė et al., 2017; Voges et al., 2019). If we consider the context of the root, *Arabidopsis* roots secrete coumarins under alkaline conditions, where iron is less bioavailable. Coumarins can both mobilize iron and act as selective antimicrobials (Voges et al., 2019). Secretion of coumarins is regulated by the transcription factor MYB72 (Stringlis et al., 2018), the expression of which can be induced by certain (coumarin-resistant) rhizobacteria (Stringlis et al., 2018). With both direct and indirect effects of iron limitation in plant/microbe communities described, identifying interactions that occur between microbes in the community can clarify the interplay between plant root and microbial community.

We had also expected to identify antibiotic compounds as active growth inhibitors in these pairwise microbe interaction assays, as our bioinformatic analysis had predicted the existence of several candidate antibiotic gene clusters in their genomes. Our functional genomics assay of *Acinetobacter* sp. 02 supernatant with the *P. putida* RB-TnSeq library did not implicate efflux pump gene clusters, which are a common resistance mechanism to protect against inhibitory small molecules (Mukhopadhyay, 2015; Eng et al., 2018). As the microbes tested in this study are representative of a synthetic model microbiome, we speculate that activation of such gene clusters would require the presence of a pathogenic or otherwise invasive microbial species. Introducing chemical or physical stressors, such as changes in temperature or humidity, DNA damaging agents, or plant hormones, could reveal new interactions between otherwise stable populations.

The methodology developed in this study enabled the examination of interactions between relevant rhizobacterial strains and revealed the role of nutrient limitation. This approach can be expanded to a much larger number of microbes and also higher order (greater than pairs) interactions. While metagenomic sequencing has identified the correlating microbial association networks present in the plant-microbe holobiont (Agler et al., 2016; Marupakula et al., 2016; Nguyen and Bruns, 2015), our study could provide the evidence to establish causal relationships which have been identified in these high throughput studies.

## Methods

### Microbial strains and cultivation

Rhizobacterial strains (Table 1) and *Pseudomonas putida* KT2440 were maintained in glycerol stocks stored at -80 °C. Microbes were routinely cultivated on lysogeny broth (LB; 10 g/L tryptone, 5 g/L yeast extract, and 5 g/L NaCl) with 10g/L agar. For nutrient-rich microbe-microbe interaction experiments, 1X LB was used with or without supplementation with 10 g/L glucose. For nutrient-poor experiments, 0.1X LB was prepared by dilution with sterile water, then supplemented with glucose (final concentration: 1 g/L tryptone, 0.5 g/L yeast extract, 0.5 g/L NaCl, 10 g/L glucose). All reagents were purchased from BD Biosciences (San Jose, CA) and were of molecular biology grade. Single colonies were obtained by streaking the desired microbe onto LB agar and incubating for 1-3 days at 30 °C, depending on the species. Liquid cultures were inoculated using single colonies from these plates. To verify taxonomy, strain identification was confirmed by analysis of 16S ribosomal sequences using the following two primers for amplification followed by Sanger sequencing: 27F: 5’-AGAGTTTGATCMTGGCTCAG-3’; 1510R: 5’-GGTTACCTTGTTACGACTT-3’. Standard protocols for PCR were followed using Q5 polymerase (New England Biolabs, Ipswitch MA). PCR was performed using manfacturer’s guidelines, and the annealing temperature was set to 50 °C for 30 cycles and a 120 second extension step at 72°C. Phylogenetic trees were generated using NCBI taxonomy IDs and visualized using iTOLv4 (Letunic and Bork, 2019).

### Bioinformatics Analysis of Candidate Secondary Metabolite Gene Clusters

Microbial genomes in this study were analyzed using bacterial antiSMASH 3.0 (Weber et al., 2015). Genomes were inputted into the algorithm using the appropriate NCBI TaxonID with the following activated parameters: default strictness; KnownClusterBlast; ActiveSiteFinder; SubClusterBlast.

### Characterization of Microbe-Microbe Interactions

For both the concurrent and staggered inoculation assays, microbe-microbe interactions were characterized based on inhibition of colony formation on an agar plate (**Figure 1**). Due to the order in which primary and secondary strains were applied to the agar plate, regions were formed for each secondary strain in which plated cells were inoculated alone and mixed with other microbial species (see **Figure 2**). Microbial growth was inspected every 24 hours for the appearance of colony forming units (CFUs) or when many viable cells were present, bacterial lawns. If both microbes showed similar CFU formation when comingled or free from the presence of a second microbe, there was no observed interaction. Where we observed apparent reductions in growth of one or both microbes, these interactions were classified as inhibitory. We further defined a category for morphological changes wherein both microbes could grow in each other’s presence, but had a change in colony or lawn formation. With few exceptions, plate based assays were repeated with >3 biological replicates over many weeks, using different batches of prepared solid agar media. Experiments were typically started using single colonies struck out from glycerol stocks no longer than 5 days prior.

### Concurrent inoculation assays

Agar (1% w/v) plates were prepared containing either 1X or 0.1X LB ± 55 mM D-glucose as previously described (Nguyen et al., 2011; Rigali et al., 2008). A representative plate is shown in Figure 2. The concentration of iron in LB was calculated using specifications provided in the “Bionutrients Technical Manual Vol 3” supplied by the manufacturer (BD Biosciences). For assays testing the role of iron availability, plates were supplemented with 1, 10, or 100 µM FeCl_3_. Single colonies of a candidate microbe, designated the primary species, were first streaked along a plate in two parallel, vertical lines. Immediately afterwards, colonies from 4-6 secondary candidate microbes were blotted at a small point roughly equidistant from each streak of the primary species. A sterile toothpick was then used to streak the secondary species in a line perpendicular to the primary species such that they intersect and mix. Plates were then incubated at 30 °C, imaged daily for 1-3 days using a Canon 550D camera, and images were saved as JPEG files for further analysis. The brightness was adjusted uniformly to maximize visibility. This experimental design allowed for observation of both direct cell-cell contact, as well as effects that happen at a distance. Antagonism between microbes was defined as a visible zone of growth inhibition in one or both species as a result of their co-culture.

### Assessment of Microbial Interactions on Spent Solid Media

A sterile 0.44 µm nitrocellulose membrane (1 × 5 cm) was applied to one half of the agar plate, prepared as described above. The primary species was inoculated vertically along the plate, with one streak on top of this membrane, and one directly onto the agar surface. Plates were incubated for 2 days at 30 °C, after which the nitrocellulose membrane was removed along with any associated bacteria. At this point, the inoculation of secondary species, imaging, and analysis proceeded as in the concurrent growth assays described above.

### Generation of Liquid Spent Culture Media

To test if a growth inhibitory compound was present in liquid culture media, *Acinetobacter* sp. 02 was inoculated into 5 mL of 0.1X LB + 55 mM D-glucose and grown for 3 days (200 RPM, 30 °C). Spent medium was obtained by centrifuging the saturated culture and filtering the supernatant at 0.22 µm. Spent media was kept at room temperature until use, which was no longer than 1 month. Either WT *P. putida* KT2440 or a *P. putida* KT2440 pooled transposon library (described in (Rand et al., 2017)) was inoculated into this spent media. Cultures were incubated at 30°C shaking at 200 RPM and samples for fitness analysis were taken 3 hours post inoculation.

### RB-TnSeq Fitness Analysis

A 1 mL aliquot of the *P. putida* RB-TnSeq library described previously was used to inoculate 25 mL of 0.1X LB + 55 mM D-glucose in a 250 mL baffled flask and grown, shaking at 200 RPM, at 30°C overnight. Spent liquid media from Acinetobacter sp.02 was generated by growing *Acinetobacter* sp.02 in 25 mL 0.1X LB + 55 mM D-glucose for 3 days in a baffled shake flask at 200 rpm. After 3 days of growth, the culture was pelleted by centrifugation at 4000 rcf for 10 min and supernatant was recovered by sterile filtration through a 0.45 µM filter. The P. putida RB-TnSeq pooled mutant library was inoculated into either 0.1X LB + 55 mM D-glucose or the spent media. Samples were taken as a “Time 0” and the preculture was used to inoculate 700 µL of 100% *Acinetobacter* sp.02 spent media or control media per well of a 48 well plate. Plates were sealed with a gas-permeable membrane and incubated at 30°C overnight, shaking at 200 RPM. Undiluted *Acinetobacter* sp. 02 spent media used here does not allow for the growth of *P. putida* KT2440; therefore after a 24 hour incubation we back-diluted the inoculated spent media 1:10 into fresh 0.1X LB + D-glucose in a 48 well plate. After an overnight outgrowth samples were pelleted and frozen at -80C for barcode sequencing. We performed DNA barcode sequencing as previously described (Wetmore et al., 2015). The fitness of a strain is defined here as the normalized log_2_ ratio of barcode reads in the experimental sample to barcode reads in the time zero sample. The fitness of a gene is defined here as the weighted average of strain fitness for insertions in the central 10% to 90% of the gene. All fitness data in this work is publicly available at http://fit.genomics.lbl.gov.

## Acknowledgements

This work was funded by a Laboratory Directed Research and Development (LDRD) grant at LBNL. TE, AM, and JM were also funded as part of the DOE Joint BioEnergy Institute (http://www.jbei.org) supported by the US Department of Energy, Office of Science, through contract DE-AC02-05CH11231 between Lawrence Berkeley National Laboratory and the US Department of Energy.

## Data Availability

The raw data supporting the conclusions of this manuscript will be made available by the authors, without undue reservation, to any qualified researcher. In particular, the collected photomicrographs of all microbial growth assays used to assess microbial interactions are available upon request. The RB-TnSeq fitness data described in Figure 5 is available for access at http://fit.genomics.lbl.gov.

## Contributions

AM, JM, TE, and RH conceived the study. RH, BW, and UM collected and analyzed the data. TE and AM helped organize the data and project. BB, JC, MB, and AD contributed valuable reagents and technical expertise. BB, JM, MB, and AM provided funding. TE and RH wrote the first draft of the manuscript and prepared figures. All authors took part in editing the manuscript and have read and approved the final version.

## Tables

**Table 1.** Summary of microbes used in this study.

## Supplemental Data Legends

**Supplemental Figure 1.**
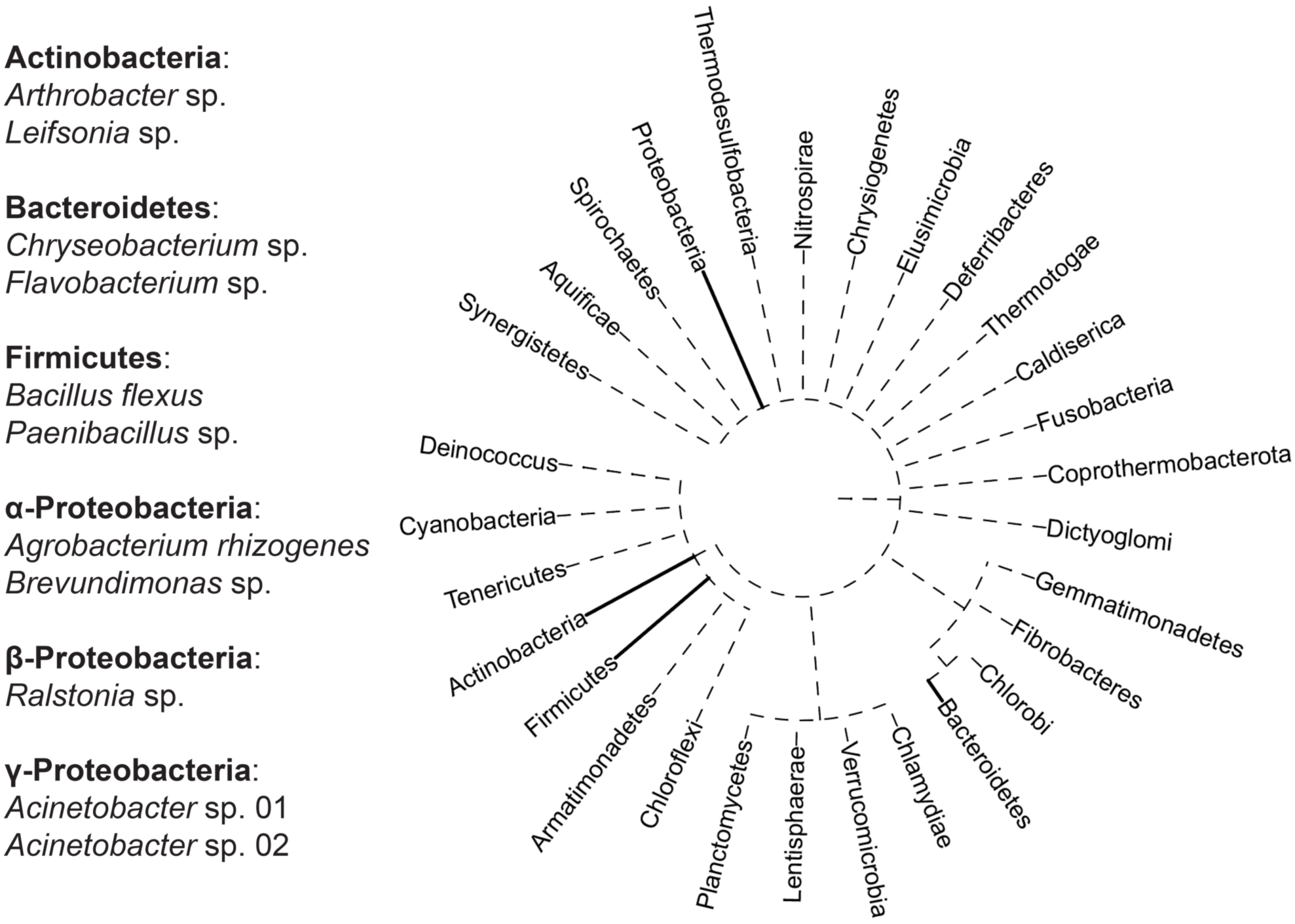
Phylogeny of rhizobacteria used in this study. Genus-level description of selected rhizobacteria and their representation within major bacterial phyla. The phylogenetic tree was generated using NCBI taxonomy IDs and visualized using iTOL V4.

**Supplemental Figure 2.**
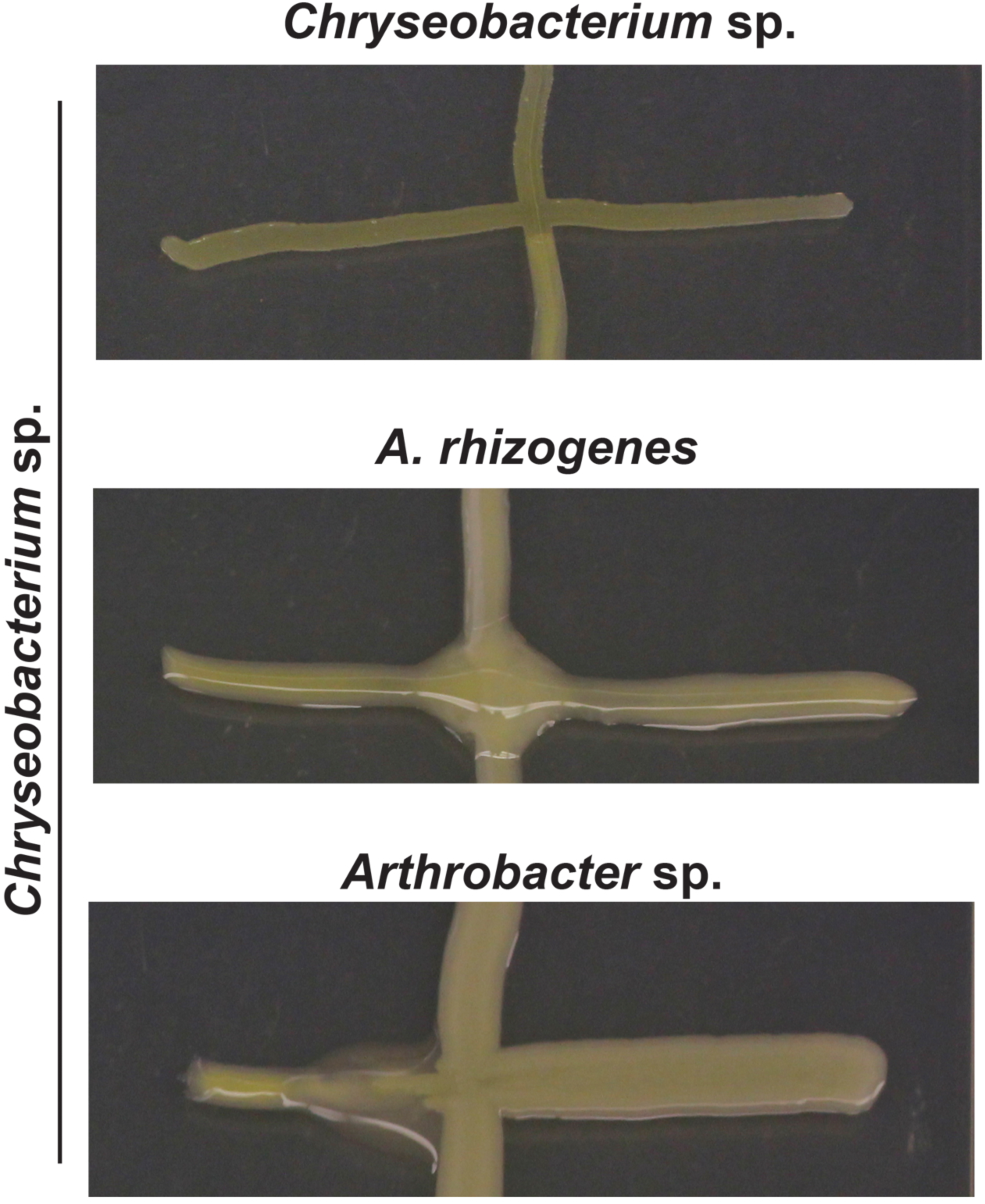
Representative images of *Chryseobacterium* sp. grown in co-culture with other rhizobacteria. *Chryseobacterium* biomass changes color and opacity when co-cultured with *A. rhizogenes* or *Arthrobacter* sp., but not when co-cultured with itself.

**Supplemental Figure 3.**
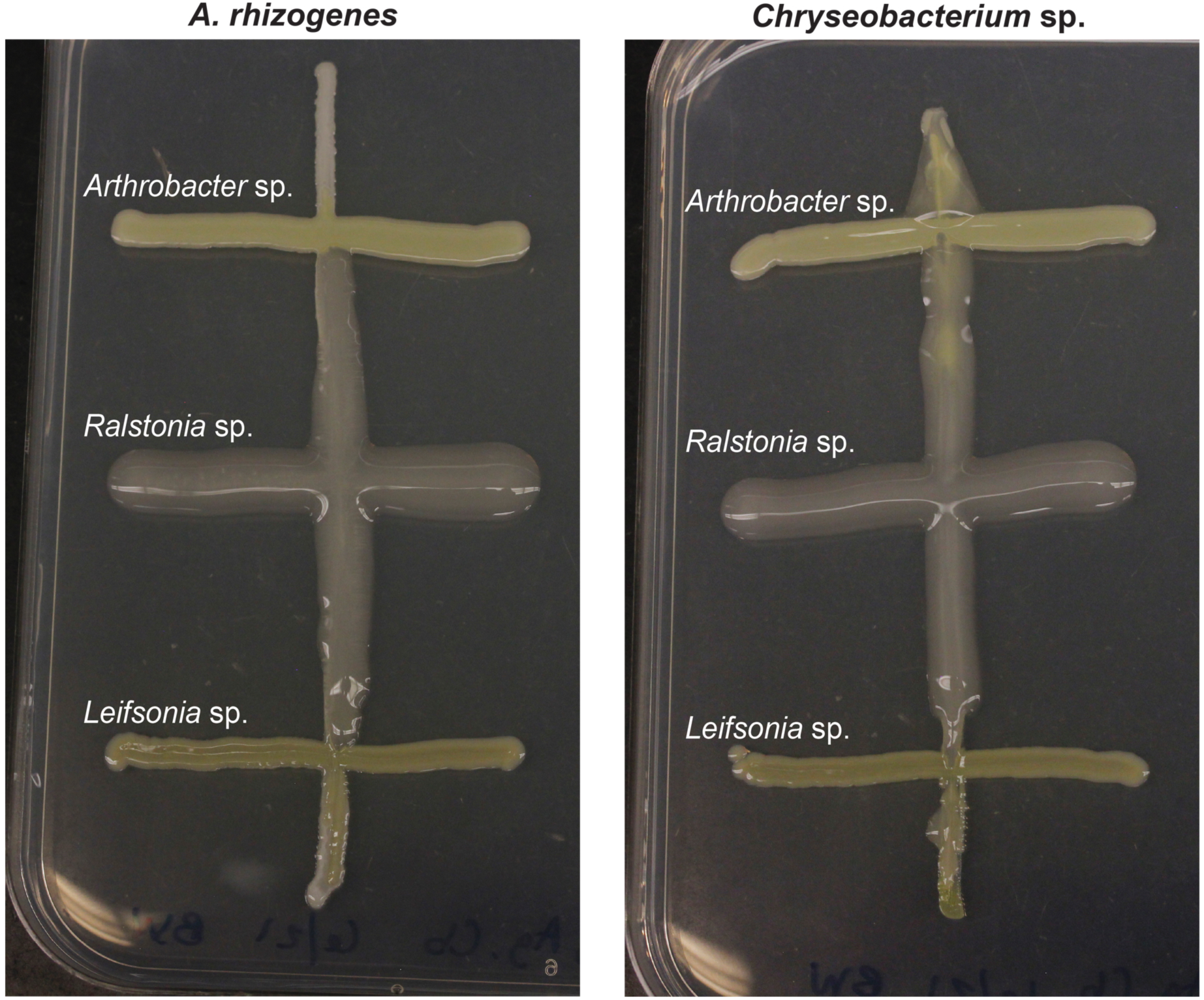
Representative images of *Ralstonia* sp. growing along the biomass of other rhizobacteria.

**Supplemental Figure 4.**
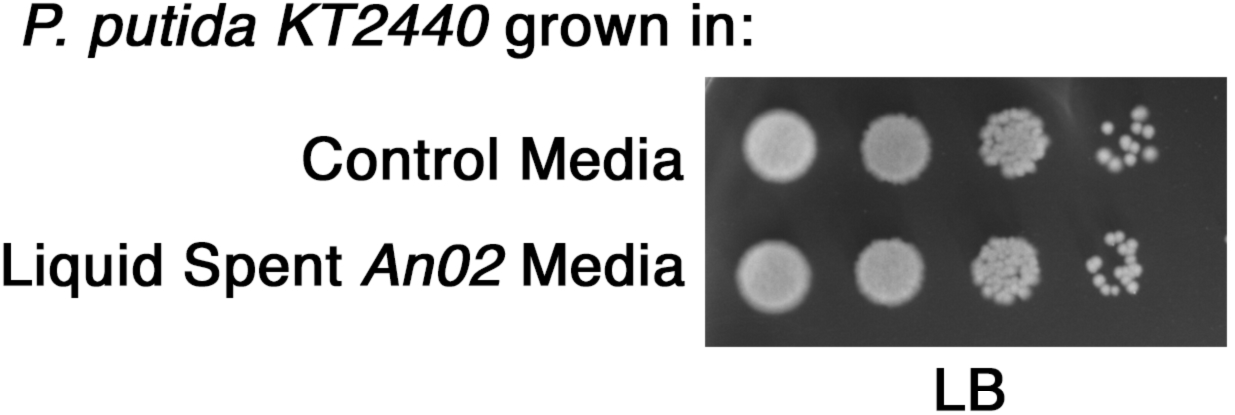
*P. putida* KT2440 grows in *Acinetobacter* sp. 02 supernatant. A log phase culture of *P. putida* KT2440 was back-diluted into fresh 1/10x LB + glucose, or spent liquid media from *Acinetobacter* sp. 02. After 3 hours of growth at 30°C, cultures were serially diluted and plated onto LB solid agar media to assess colony formation.

**Supplemental Figure 4.**
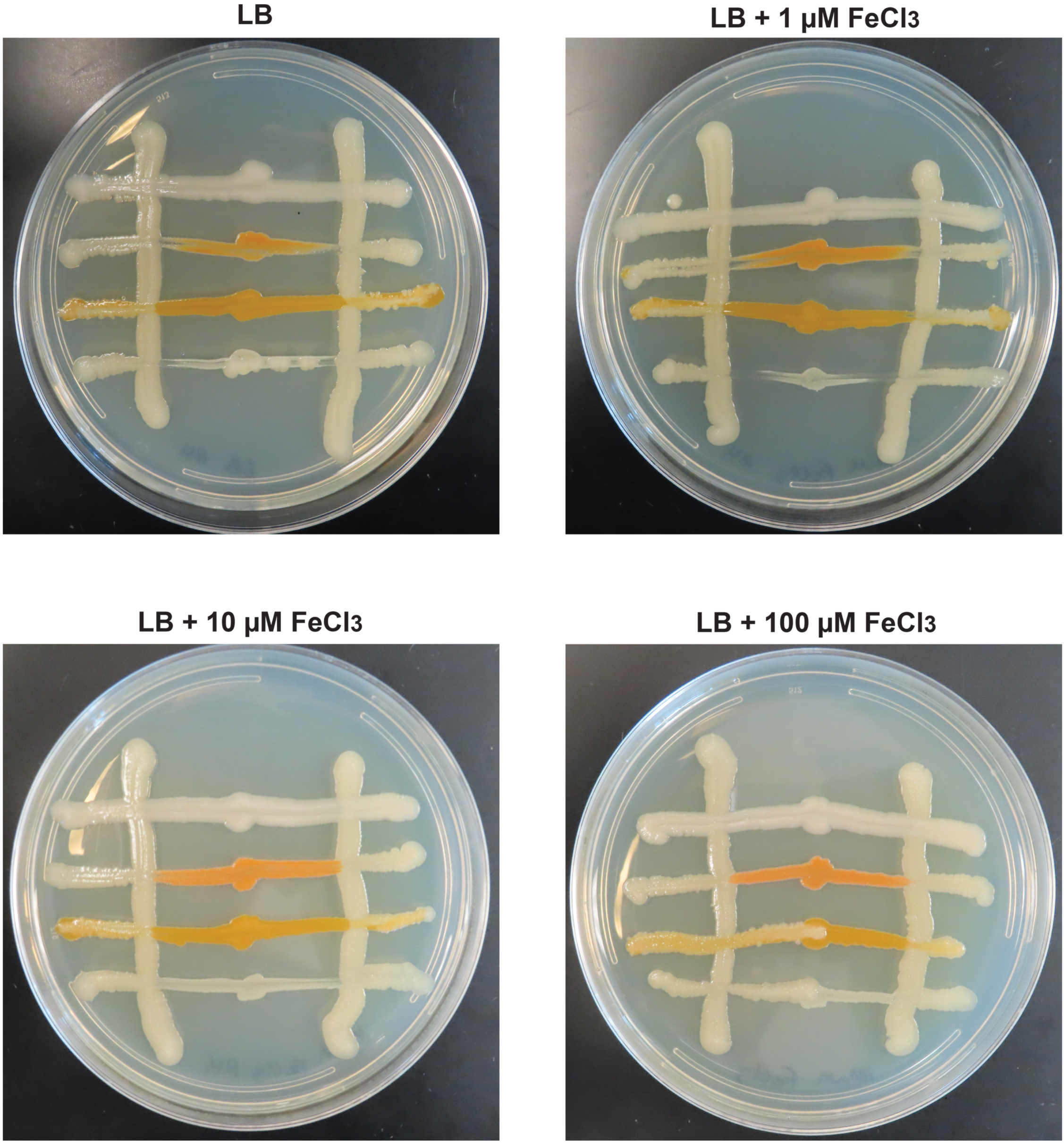
The effect of iron on interactions between *Acinetobacter* sp. 02 and other rhizobacteria is dose dependent. N ≥ 3.

**Supplemental Table 1.** Summary of anti-SMASH gene clusters identified from microbes in this study.

