## Supplementary material for "Iron Supplementation Eliminates Antagonistic Interactions Between Root Associated Bacteria": Table 1

| **Microbe** | **NCBI Taxonomy ID** | **JBEI Accession number** |
| --- | --- | --- |
| 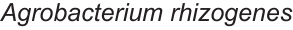 | 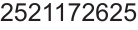 | **JBEI-16051*** |
| 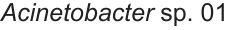 | 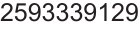 | **JBEI-16052** |
| 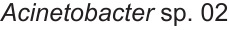 | 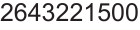 | **JBEI-16055*** |
| 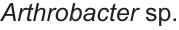 | 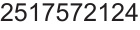 | **JBEI-16049** |
| 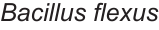 | 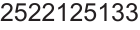 | JBEI-16054 |
| 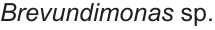 | 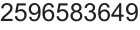 | JBEI-16062 |
| 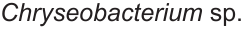 | 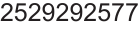 | JBEI-16083 |
| 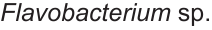 | 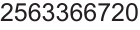 | JBEI-16082 |
| 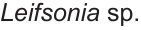 | 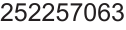 | JBEI-16050 |
| 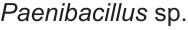 | 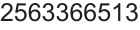 | JBEI-16053 |
| 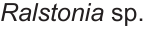 | 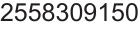 | JBEI-16059 |

*The description of these strains has been adjusted from their original NCBI taxonomy following full-length pyrosequencing of the bacterial 16S gene using universal 27F and 1492R primers.
